# Thermodynamic Constraints on Electromicrobial Protein Production

**DOI:** 10.1101/2021.11.22.469619

**Authors:** Lucas Wise, Sabrina Marecos, Katie Randolph, Eric Nshimyumukiza, Mohamed Hassan, Jacob Strouse, Farshid Salimijazi, Buz Barstow

## Abstract

Global consumption of protein is projected to double by the middle of the 21^st^ century^1^. However, protein production is one of the most energy intensive and environmentally damaging parts of the food supply system today. Electromicrobial production technologies that combine renewable electricity and CO_2_-fixing microbial metabolism could dramatically increase the energy efficiency of commodity chemical production^2–5^. Here we present a molecular-scale model that sets an upper limit on the performance of any organism performing electromicrobial protein production. We show that engineered microbes that fix CO_2_ and N_2_ using reducing equivalents produced by H_2_-oxidation or extracellular electron uptake could produce amino acids with energy inputs as low as 64 MJ kg^-1^. This work provides a roadmap for development of engineered microbes that could significantly expand access to proteins produced with a low environmental footprint.

## Introduction

### Current methods of protein production are environmentally damaging

Current food consumption and farming practices produce a large amount of environmental strain. In particular, the production of livestock for protein leads to significant waste accumulation and energy expenditure^6^. The agricultural and food production sectors are responsible for ≈ 30% of greenhouse gas emissions, while livestock farming alone accounts for 18% of emissions^7^. Furthermore, the agricultural industry is responsible for 70% of total freshwater consumption^8^. 42% of freshwater consumption is attributed to livestock production alone^8^. But, increased consumption of protein is one of the best ways to improve human, particularly infant, health and productivity in many parts of the world today^9^.

The energy and water consumption of livestock farming will only increase as global appetites increase. First, population will grow to ≈ 11 billion by 2050^10^. Second, the consumption of food, particularly protein, by each individual will also grow thanks to an expected average annual economic growth rate of 3% from 2014 to 2050^11,12^. Supplying this increased demand while maintaining the current agricultural areal footprint is expected to require a 75% increase in agricultural productivity^10^.

Should agricultural production efficiencies remain stagnant, satisfying the food demands of the world’s growing and increasingly wealthy population with protein will require massive deforestation^13,14^. Deforestation could eradicate thousands of species and produce large quantities of greenhouse gases, leading to temperature increases exceeding the 2 °C warming threshold established by the Paris Climate Agreement, even when ignoring emissions from all other human activity^15^.

Incremental improvements in current food production technologies may not meet future demand and sustainability goals. Current approaches to increasing protein production include advanced livestock breeding, and substitution of livestock protein for insect- and plant-based substitutes. However, all of these approaches depend upon increases in crop yields. But, 78% of the world’s land has natural limitations for agricultural development^10^, and significant doubts remain about the possibility of increasing crop yields by mid-century^11,16,17^. Furthermore, increasing water scarcity due to climate change could even depress crop yields in the decades ahead^16^.

### Autotrophic metabolism could increase the efficiency of protein production

Autotrophic microbial production of protein is a promising alternative strategy to conventional food production^5,18–20^. In this class of schemes, externally supplied reducing equivalents are used to power microbial N_2_ and CO_2_-fixing metabolism and synthesis of protein molecules^21,22^.

In most systems studied to date, reducing equivalents are supplied by H_2_^-^ or methane-oxidation. CO_2_-fixation is performed by Calvin-Benson-Bassham cycle, the reverse Krebs cycle or the Wood-Ljungdahl pathway.

Autotrophically produced protein has at least three important advantages over traditional protein production methods. First, autotrophic microorganisms can already produce protein at much higher rates than traditional protein sources^18,19^. Secondly, autotrophic protein production does not depend on the availability of arable land and can be run in a closed system. This greatly reduces water and land consumption and inhibits nitrogen runoff to surrounding environments^19,23^. Finally, autotrophic microorganisms can use atmospheric N_2_ as a substrate, eliminating the need for thermochemical N_2_-fixation^18,19^.

The cost of autotrophic protein production is dropping rapidly. The cost of production of a single protein has reduced from $1 × 10^6^ kg^-1^ in 2000 to ≈ $100 kg^-1^ in 2019^24^. It is projected that the cost of production of a single protein could drop to below $10 kg^-1^ by 2025, thereby achieving price parity with animal-based protein products^24^.

Theoretical analysis suggests that autotrophic protein production could far exceed the efficiency of plant-based protein. Recent analyses of the performance of electromicrobial production of biofuels^2,4^, where electrically-supplied reducing equivalents are used to power CO_2_ fixation or formic acid assimilation and biofuel, show that these types of schemes could dramatically exceed the efficiency of photosynthetic biofuel production. These results imply that if N_2_ fixation were added to these systems, proteins could also be produced at efficiencies exceeding that of photosynthesis. Recent results by Leger *et al*.^5^ suggest photovoltaic-driven electromicrobial production (EMP) of protein could exceed efficiency of real-world photosynthetic production of protein by at least 2 orders of magnitude.

However, up until now, very few attempts have been made at calculating the upper limit efficiency of EMP amino acid or protein production. This paper presents a model and analyzes the theoretical maximum energetic efficiency for a system of autotrophic microorganisms, fixing CO_2_ and N_2_ using electrons delivered by either extracellular electron uptake (EEU)^25^ or by H_2_-oxidation^26^. These calculations do not predict the performance of any naturally-occurring organism, but do predict an upper limit efficiency for any natural or synthetic organism using these reactions.

## Theory, Results and Discussion

### Theory

We extended our theoretical framework for calculating the efficiency of electromicrobial production (EMP) of biofuels to calculate the efficiency of amino acid production from electrons, CO_2_ and N_2_ (ref. ^4^). A full set of model parameters and associated values used in this article are shown in **Table 1**, and a full set of symbols for this article are shown in **Table S1**.

**Table 1.**
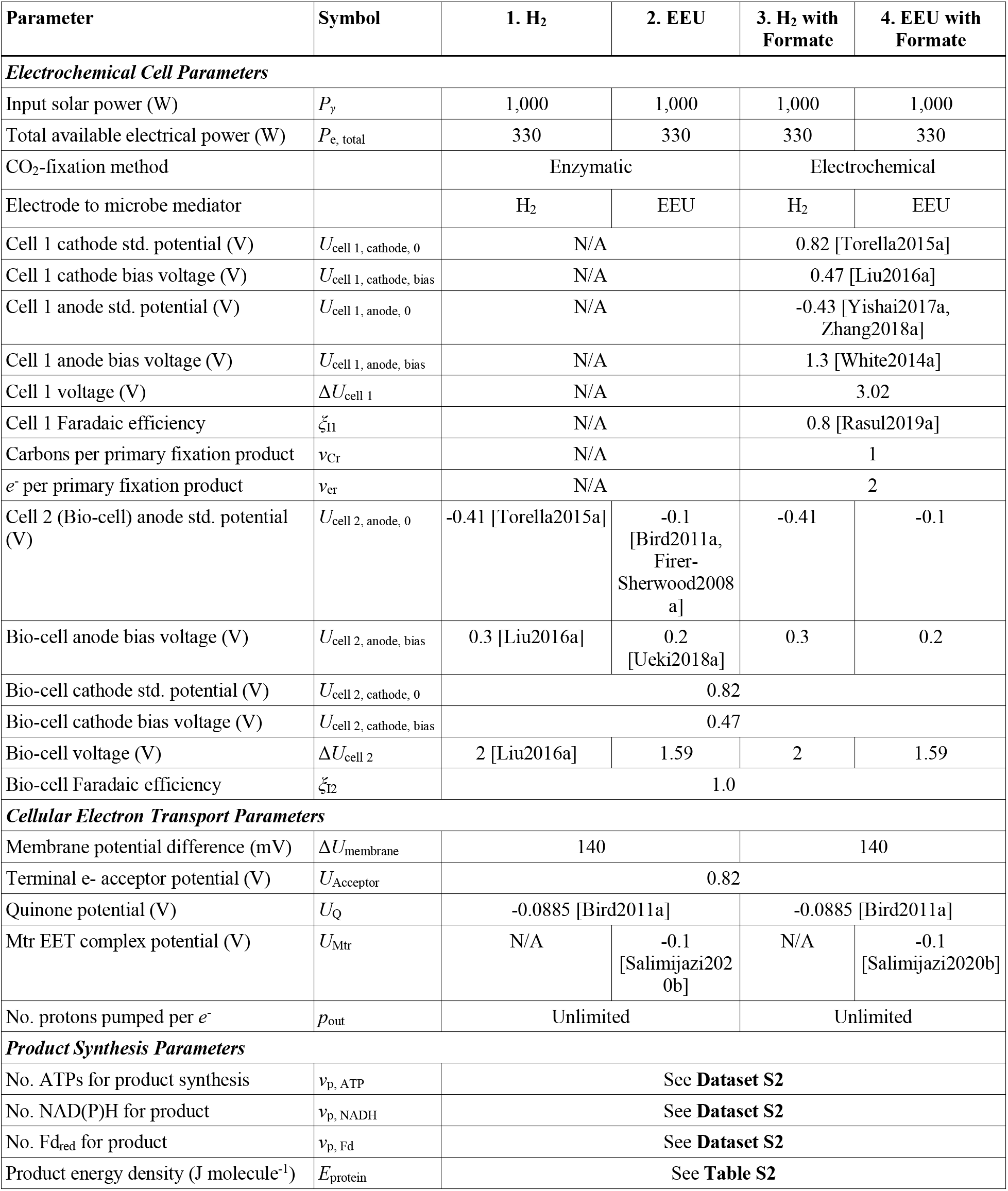
Electromicrobial protein production model parameters. Model parameters used in this article are based upon model parameters used in a previous analysis of the electromicrobial production of the biofuel butanol^4^. A sensitivity analysis was performed for all key parameters in this work^4^.

We consider a bio-electrochemical system used to deliver electrons to microbial metabolism (**Figures 1A** and **1B**). Electrical power is used to generate amino acid (or protein) molecules with an energy per molecule *E*_protein_ at a rate *Ṅ*_protein_. Even though this article strictly considers amino acid synthesis, this can be considered equivalent to protein production from an energetic standpoint as no energy is expended in forming the peptide bond needed to polymerize amino acids. We choose to use the subscript protein rather than AA to avoid confusion with the Avogadro constant, *N*_A_. Energy per molecule and molecular weight for each amino acid are shown in **Table S2**.

**Figure 1.**
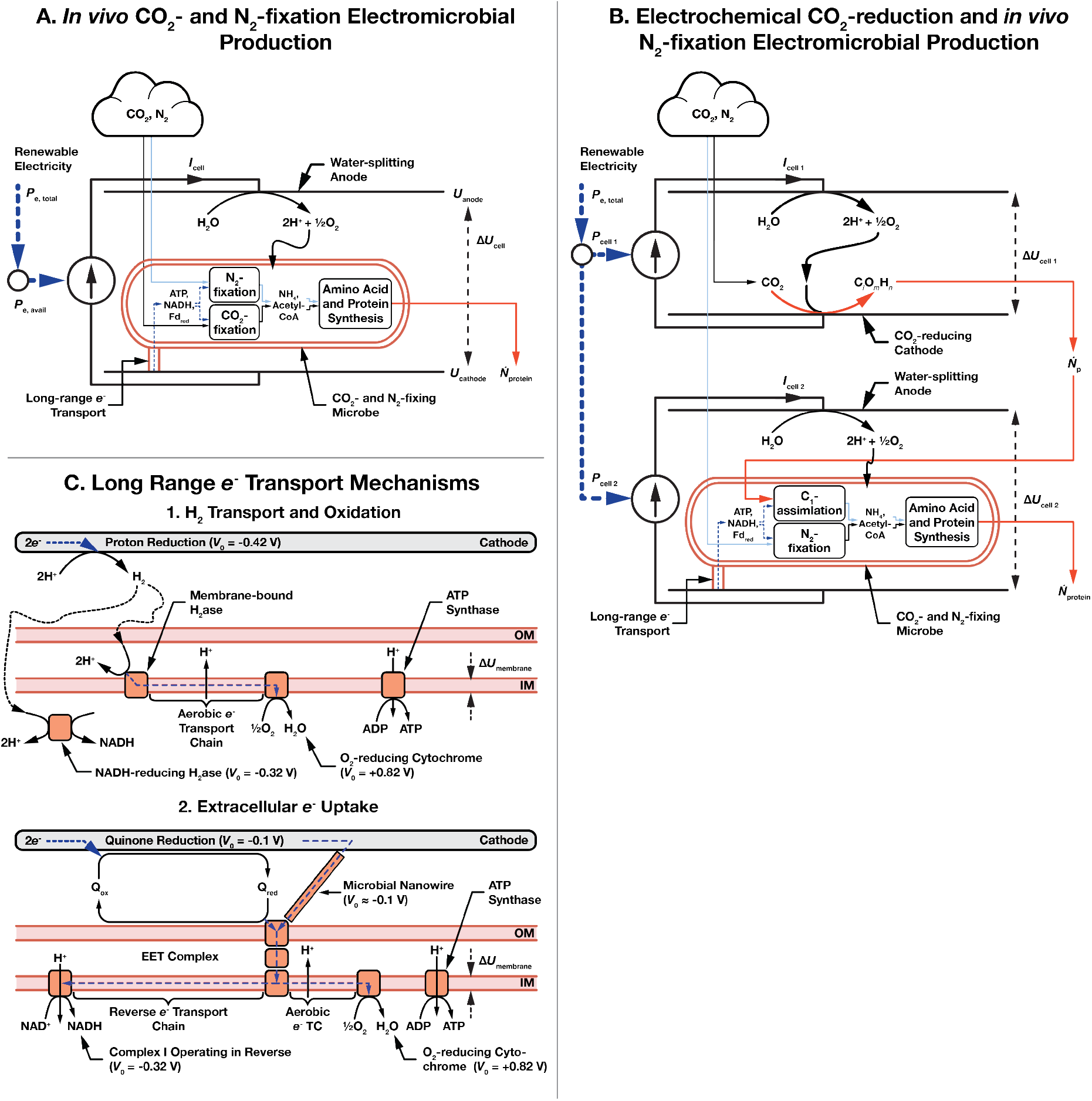
Schematic of amino acid electromicrobial production systems. (**A**) Single bio-electrochemical cell system where electricity is used to power *in vivo* CO_2_- and N_2_-fixation. (**B**) Dual electrochemical cell system where CO_2_ is reduced in the first cell, and then assimilated in the second cell, and combined with enzymatically fixed N_2_. (**C**) Long range *e^-^* transfer mechanisms considered in this article. In the first, H_2_ is electrochemically reduced on a cathode, transferred to the microbe by diffusion or stirring, and is enzymatically oxidized. In the second mechanism, extracellular electron uptake (EEU), *e^-^* are transferred along a microbial nanowire (part of a conductive biofilm), or by a reduced medium potential redox shuttle like a quinone or flavin, and are received at the cell surface by the extracellular electron transfer (EET) complex. From the thermodynamic perspective considered in this article, these mechanisms are equivalent. Electrons are then transported to the inner membrane where reverse electron transport is used to regenerate NAD(P)H, reduced Ferredoxin (not shown), and ATP^25,32^.

The energy conversion efficiency of the system from electricity to amino acids (or protein) is calculated from the ratio of the amount of chemical energy stored per second (*Ṅ*_protein_ *E*_protein_), relative to the power input to the system, *P*_e, total_ (ref. ^4^),

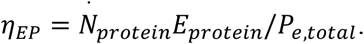

The amount of electricity needed to produce a unit-mass of the protein is,

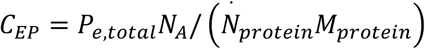

where *M*_protein_ is the molecular weight of the protein molecule.

For a single bio-electrochemical cell system where CO_2_- and N_2_- fixation are performed *in vivo* (**Figure 1A**), the upper limit electrical to chemical conversion efficiency of the system is set by the energy density of an amino acid molecule relative to the amount of charge needed to synthesize it from CO_2_ and N_2_ (the fundamental charge, *e*, multiplied by the number of electrons needed for synthesis, *ν*_ep_) and the potential difference across the bio-electrochemical cell, Δ*U*_cell_,

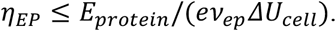

Thus, the amount of electricity needed to produce a unit-mass of the protein is,

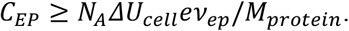

For systems where CO_2_ reduction is performed electrochemically, and the resulting reduction product (typically a C_1_ compound like formic acid) (refs. ^27–29^) is further reduced enzymatically (**Figure 1B**), *ν*_ep_ is substituted for number of electrons needed to convert the C_1_ product into the final protein product, *ν*_e, add_ (ref. ^4^),

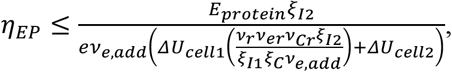

where *ν*_r_ is the number of primary reduction products (*i.e*., formic acid molecules) needed to synthesize a molecule of the final product, *ν*_er_ is the number of electrons needed to reduce CO_2_ to a primary reduction product (*i.e*., 2 in the case of formic acid), *ν*_Cr_ is the number of carbon atoms per primary fixation product (*i.e*., 1 in the case of formic acid), *ξ*_I2_ is the Faradaic efficiency of the bio-electrochemical cell, *ξ*_I1_ is the Faradaic efficiency of the primary abiotic cell 1, *ξ*_C_ is the carbon transfer efficiency from cell 1 to cell 2.

Thus, the amount of electricity needed to produce a unit-mass of the protein when using electrochemical CO_2_-reduction is,

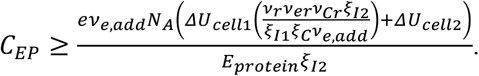

We calculate the electron requirements, *ν*_ep_ or *ν*_e, add_, for amino acid (or protein) synthesis from the number of NAD(P)H (*ν*_p, NADH_) reduced Ferredoxin (Fd_red_; *ν*_p, Fd_) and ATP (*ν*_p, ATP_) molecules needed for the synthesis of the molecule, along with a model of the mechanism used for electron delivery to the microbe^4^.

For systems that rely upon H_2_-oxidation for electron delivery like the Bionic Leaf^4,26,30^ (**Figure 1C** part 1),

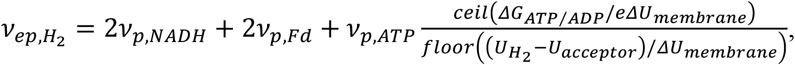

where Δ*G*_ATP/ADP_ is the free energy required for regeneration of ATP, Δ*U*_membrane_ is the potential difference across the cell’s inner membrane due to the proton gradient, *U*_H2_ is the standard potential of proton reduction to H_2_, *U*_acceptor_ is the standard potential of terminal electron acceptor reduction (typically O_2_ + 2*e^-^* to H_2_O), the ceil function rounds up the nearest integer, and the floor function rounds down to the nearest integer.

The inner membrane potential difference, Δ*U*_membrane_, is the largest source of uncertainty in this calculation. Therefore, we present a range of efficiency estimates in **Figures 2** and **3** and throughout the text for Δ*U*_membrane_ = 80 mV (BioNumber ID (BNID) 10408284^31^) to 270 mV (BNID 107135), with a central value of 140 mV (BNIDs 109774, 103386, and 109775).

**Figure 2.**
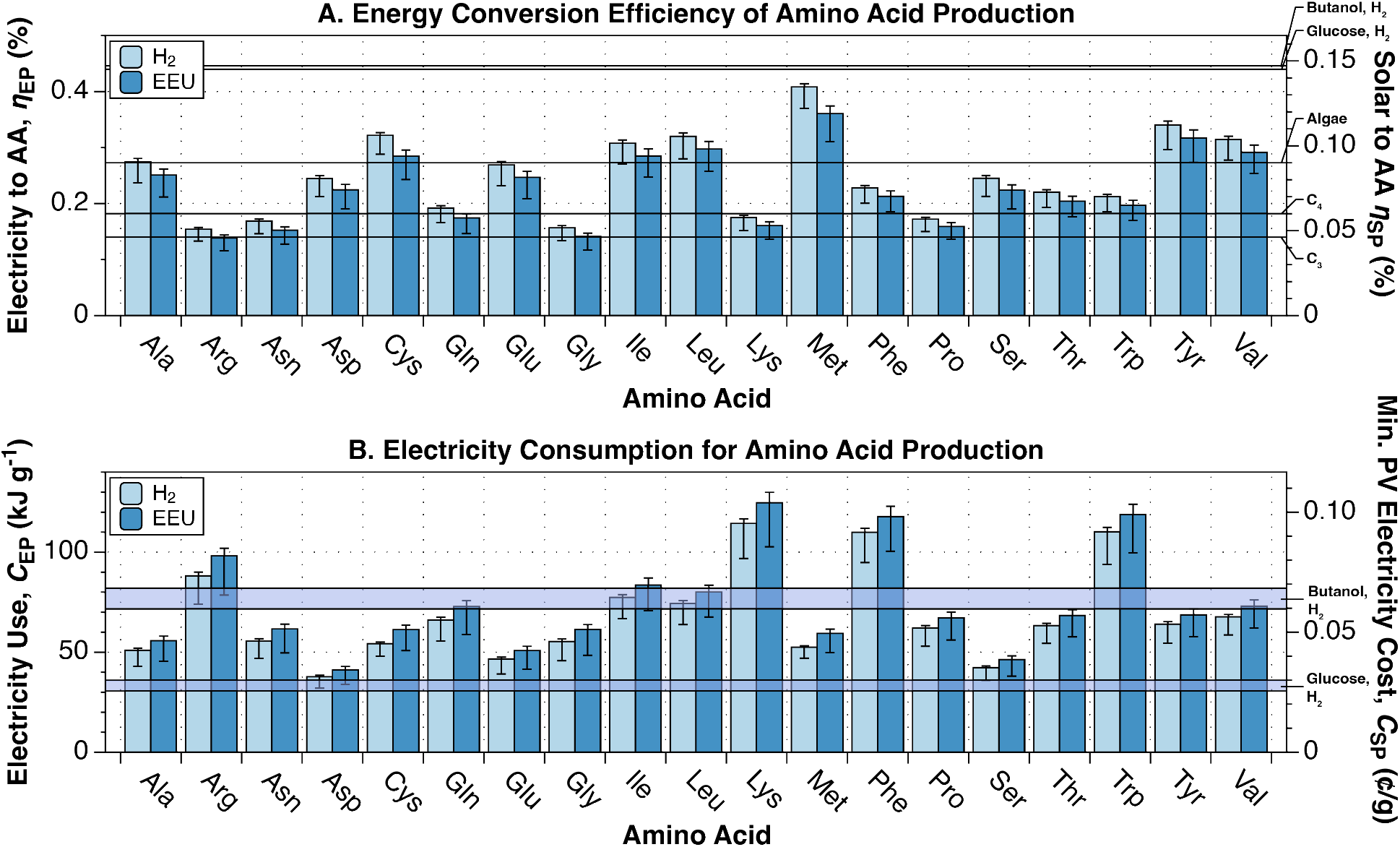
Energy conversion efficiency and energy cost of amino acid production. The upper limit energy conversion efficiency and minimum energy cost of amino acid production from CO_2_, N_2_ and electricity by electromicrobial production systems using the Calvin cycle for CO_2_-fixation and either H_2_-oxidation or extracellular electron uptake (EEU) were calculated for 19 dietary amino acids (all except histidine) with the electrofoods package^43^. NADH, Fd_red_, and ATP requirements for synthesis of each amino acid are tabulated in **Dataset S2**. This plot can be reproduced using the fig-cbb_n2_to_amino_acids.py program in the electrofoods package^43^. (**A**) Upper limit electrical and solar energy conversion efficiency for amino acids. The left axis shows the electricity to amino acid energy conversion efficiency, while the right axis shows the solar to amino acid conversion efficiency, assuming the system is supplied by a perfectly efficient single-junction Si solar photovoltaic (solar to electrical efficiency of 32.9% (ref. ^69^)). As a first point of comparison, the upper limit solar to biomass energy conversion efficiencies of C_3_, C_4_ (refs. ^44,45^), and algal photosynthesis^46^ are marked on the right axis. As a second point of comparison, we have also marked the projected upper limit solar to butanol^4^ and glucose (calculated here) conversion efficiencies by an electromicrobial production system using H_2_-oxidation and the Calvin cycle. (**B**) Minimum electrical and solar energy costs for the production of a gram of amino acids. The left axis shows the minimum electricity cost, while the right axis shows the minimum cost of that solar electricity, assuming that the US Department of Energy’s cost target of 3 ¢ per kWh by 2030 can be achieved^47^.

**Figure 3.**
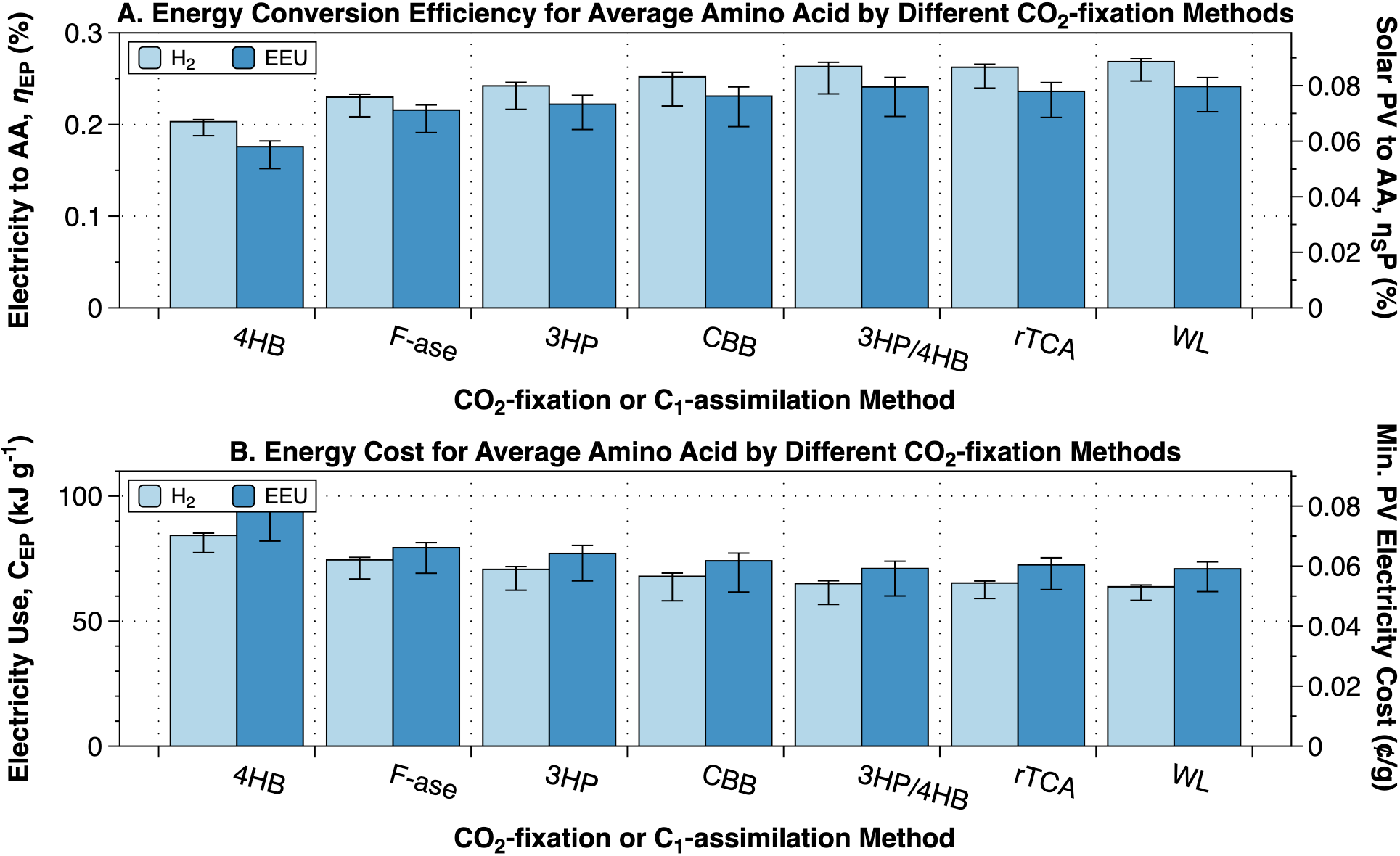
Changing CO_2_-fixation method can improve the performance of amino acid synthesis. The upper limit energy conversion efficiency and minimum energy cost of production of an average amino acid from CO_2_ or HCOO^-^, N_2_ and electricity by electromicrobial production systems using either H_2_-oxidation or extracellular electron uptake (EEU) and one of the 6 naturally-occurring CO_2_-fixation pathways or the synthetic Formolase formate assimilation pathway were calculated with the electrofoods package^43^. NADH, Fd_red_, and ATP requirements for synthesis of an average amino acid are tabulated in **Dataset S2**. This plot can be reproduced using the fig-cbb_n2_to_amino_acids.py program in the electrofoods package^43^. 3HP: 3-hydroxypropionate cycle; 3HP-4HB: 3-hydroxypropionate/4-hydroxybutyate cycle; 4HB: 4-hydroxybutyate cycle; CBB: Calvin-Bensson-Bassham cycle; Form: Formolase pathway; rTCA: reductive TCA cycle; WL: Wood-Ljungdahl pathway. (**A**) Upper limit electrical and solar energy conversion efficiency for an average amino acid. The left axis shows the electricity to amino acid energy conversion efficiency, while the right axis shows the solar to amino acid conversion efficiency, assuming the system is supplied by a perfectly efficient single-junction Si solar photovoltaic (solar to electrical efficiency of 32.9%^69^). (**B**) Minimum electrical and solar energy costs for the production of a gram of an average amino acid. The left axis shows the minimum electricity cost, while the right axis shows the minimum cost of that solar electricity, assuming that the US Department of Energy’s cost target of 3 ¢ per kWh by 2030 can be achieved^47^.

For systems that rely upon EEU for electron delivery like *Shewanella oneidensis*^4,25^ (**Figure 1C** part 2),

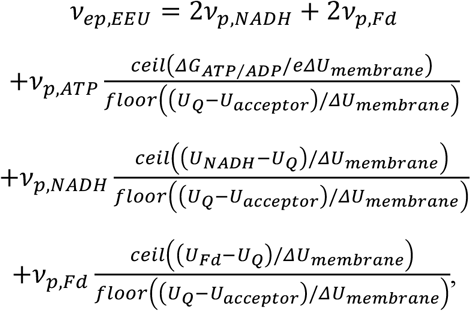

where *U*_Q_ is the redox potential of the inner membrane electron carrier, thought to be ubiquinone^32^, *U*_NADH_ is the standard potential of NAD(P)H, and *U*_Fd_ is the standard potential of Ferredoxin.

The NAD(P)H, ATP and Fd_red_ requirements for amino acid synthesis were calculated by balancing networks of reactions for the autotrophic synthesis of the molecule from N_2_ and CO_2_ or N_2_ and formate (COOH^-^). We enumerated all reaction steps for the production of 19 of the 20 dietary amino acids from acetyl-CoA and NH_4_ using data from the KEGG database in **Dataset S3** (refs. ^33–35^). Synthesis of histidine was excluded from these calculations because of technical challenges with stoichiometric balancing due to its inseparable connection with purine synthesis. As a comparison point, and to validate our approach, we also consider the synthesis of glucose.

Amino acid synthesis reactions were complemented with reactions for CO_2_-fixation, C_1_-assimilation, and N_2_ fixation (**Table S3**). For this article we considered 6 scenarios in which CO_2_ was fixed by the well-known Calvin cycle^36^, the Reductive Tricarboxylic Acid cycle^37,38^, Wood-Ljungdahl (WL) Pathway^36^; the 3-hydroxypropionate/4-hydroxybutyrate (3HP-4HB) Pathway^38,39^; 3-hydroxypropionate (3HP) Cycle^40^; and the Dicarboxylate/4-hydroxybutyrate (4HB) Cycle^41^. In addition, we also considered the artificial Formolase formate assimilation pathway^42^. Finally, in all scenarios, N_2_ was fixed into metabolism by the iron-molybdenum (FeMo) nitrogenase (Kyoto Encyclopaedia of Genes and Genomes (KEGG) reaction R05185 (refs. 22-24).

The overall stoichiometry of autotrophic amino acid synthesis was calculated by a custom flux balance code. Amino acid synthesis reactions (**Dataset S1**) were combined automatically with the CO_2_-fixation, C_1_-assimilation, and N_2_ fixation reactions (**Table S3**) by a custom code^43^ into a set of stoichiometric matrices, **S_p_**, for each reaction network.

Each automatically generated stoichiometric matrix was balanced with a custom flux balance program^43^ to find the overall number of NAD(P)H, Fd_red_, and ATP needed for synthesis of each amino acid using each CO_2_-fixation or C_1_-assimilation pathway.

We consider a species number rate of change vector, ***ṅ***, that encodes the rate of change of number of the reactant molecules over a single cycle of the reaction network; a stoichiometric matrix **S_p_** that encodes the number of reactants made or consumed in every reaction in the network; and a flux vector ***ν*** that encodes the number of times each reaction is used in the network. Reactant molecules are denoted as inputs (*e.g*., CO_2_, N_2_, COOH^-^, ATP, NAD(P)H), outputs (*e.g*., H_2_O), intermediates, or the target molecule (*e.g*., the amino acid to be synthesized). For the purposes of this thermodynamic analysis, we consider NADH and NADPH to be equivalent as they have near identical redox potentials.

The reactant number vector elements for the inputs were calculated by numerically solving the flux balance equation,

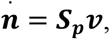

under the constraint that number of each intermediate does not change over a reaction cycle, and that number of target molecules increases by 1,

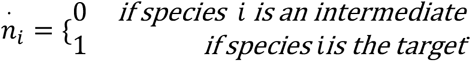

The balanced overall stoichiometry for synthesis of each amino acid is shown in **Dataset S2**.

The number of electrons needed to synthesize an average amino acid was found by calculating the average number of NAD(P)H, Fd_red_, and ATP needed for synthesizing 19 of the 20 amino acids.

### Results and Discussion

#### Electromicrobial Production of Amino Acids and Protein

The electrical and solar energy to protein conversion efficiency (*η*_EP_ and *η*_SP_) and the electrical energy consumption per unit mass (*C*_EP_) and cost of solar electricity per unit mass (*C*_SP_) for the production of 19 amino acids was calculated for electron uptake by H_2_ transport and oxidation and EEU, and CO_2_ fixation by the Calvin cycle (**Figure 2**).

Amino acid synthesis has a lower conversion efficiency than purely carbon-containing products due to the high Fd_red_ and ATP requirements of N_2_-fixation (**Dataset S1**). Despite this, the conversion efficiency either matches, and in most cases exceeds the theoretical maximum conversion efficiency of sunlight to carbohydrate biomass by C_3_ photosynthesis (**Figure 2A**). However, Arg, Asn, Gly, and Pro synthesis by H_2_ and EEU, and Gln synthesis by EEU have lower conversion efficiencies than C_4_ carbohydrate photosynthesis^44,45^. Synthesis of Cys, Ile, Leu, Met, Phe, Tyr and Val exceed the theoretical efficiency of algal photosynthesis^46^. The average CO_2_, N_2_, and electricity conversion efficiency for an average amino acid using the Calvin cycle is 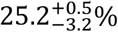when using H_2_-oxidation, and 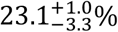when using EEU (**Figure 3A**).

The electrical energy costs (*C*_EP_) for individual amino acids using H_2_-oxidation an the Calvin cycle range from 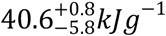 for Asp to 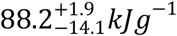 for Arg (**Figure 2B**). Synthesizing the amino acids by EEU rather than H_2_ adds between ≈ 5 and 10 kJ g^-1^. At projected 2030 prices for solar photovoltaic electricity from the DOE’s SunShot program of 3 ¢ per kWh^47^, this corresponds to a minimum cost of 0.033 ¢ g^-1^ to 0.081 ¢ g^-1^ (**Figure 2B**). The average amino acid synthesis energy cost using H_2_-oxidation and the Calvin cycle is 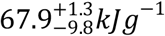 (**Figure 3B**).

As noted before, the energy conversion efficiency of systems using EEU is consistently a few percentage points lower than for systems using H_2_ oxidation^4^ (**Figure 2A**). In EEU based systems there is a higher electron requirement, and hence cell current, needed for regeneration of NAD(P)H, Fd_red_ and ATP. Practically, this is almost offset by a lower minimum cell voltage, resulting in a slightly lower conversion efficiency^4^. Averaged across all amino acids, the efficiency of synthesis for systems using EEU and the Calvin cycle is 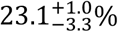. This results in an average electrical energy cost that of 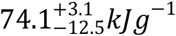, about 6*kjg*^−1^ higher than the cost of synthesis using H_2_-oxidation.

Can we use biological engineering to increase the energetic efficiency of electromicrobial production of amino acids? As we have examined before^4^, we can improve efficiency by swapping the Calvin cycle for the an alternative CO_2_ fixation cycle (**Figure 3**). As an aside, the only alternative N_2_-fixation pathway uses the iron-vanadium nitrogenase, that requires 40 ATP and 12 Fd_red_ for each N_2_ fixed (KEGG reaction R12084), compared with 16 ATP and 8 Fd_red_ for the more common iron-molybdenum-cobalt nitrogenase (KEGG reaction R05185).

Not unexpectedly, the order of efficiency of amino acid synthesis efficiency is approximately the same as the order of efficiency of butanol synthesis. As before^4^, the 4HB cycle, which performed least well for butanol synthesis^4^, also performed least well for amino acid synthesis. Likewise, the Wood-Ljungdahl pathway performed the best (**Figure 3A**).

With increasing efficiency comes decreasing electricity cost (**Figure 3B**). The average cost of producing a gram of amino acid with H_2_-4HB is 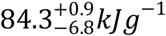 and 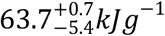 with H_2_-WL (costs of 0.07 and 0.05 ¢ g^-1^). Swapping to EEU-4HB increases the costs 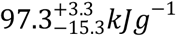, and swapping to EEU-WL reduces them to 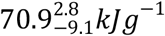 (costs of 0.08 and 0.06 ¢ g^-1^).

### Electromicrobial Protein is an energy-efficient alternative to current protein production technologies

How do the upper-limit efficiencies predicted for EMP protein production compare with real world production efficiencies and energy costs? Most rigorous estimates of the total cradle-to-farm gate energy costs needed to produce a gram of beef, chicken, pork, eggs, and dairy^48^; soybeans^49^; insects^50^ and cultured meat^14^ consider only primary energy inputs. Estimates of primary energy input start at 44 kJ g^-1^ for soybeans^49^ and go up to 273 kJ g^-1^ for beef^48^ (**Table S4**).

However, traditional estimates of energy input into protein production are not suitable for an apples-to-apples comparison to the numbers calculated in this article. These estimates consider the energy content of feed stocks such as grain and milk; and infrastructural costs such as transportation to the farm gate and tilling land. In the case of soy bean production, the estimates do not include the energy delivered by sunlight to the system to initially fix CO_2_, N_2_ and synthesize amino acids. Likewise, for livestock and dairy production, they do not include the energy content of the sunlight needed to produce the feed, only its final energy content.

Traditional energy input estimates of protein production are not wrong. Quite rightly, sunlight has been thought of as free of cost and global warming concerns. Furthermore, traditional analyses rightly concern themselves with necessary fossil energy inputs. However, as global agricultural production expands, the land for agriculture becomes an increasingly precious commodity. As a result, efficiency of use of sunlight becomes increasingly important.

Likewise, our analysis explicitly ignores infrastructural costs. While we would like to think that bioreactor production of protein could avoid many of these costs, simply thinking this does not make it so. We cannot say so with any certainty if the infrastructure energy costs, such as stirring, heating, gas exchange, are less than the energy inputs associated with agriculture or livestock farming needed to produce a gram of protein.

Estimates of photosynthetic cost of producing protein are the closest comparison point to our work. The closest comparison point to this work is a recent comparison of year round production of protein rich crops, and their protein content with an empirical model of electromicrobial production methods by Leger *et al*.^5^. The analysis by Leger *et al*. allows for calculation of the solar energy costs of photosynthetic production (**Table S5**). Energy costs range from 47 MJ g^-1^ (*η*_SP_ = 0.035%) for soybeans grown in the US to 408 MJ g^-1^ (*η*_SP_ = 0.004%) for maize grown in India (**Table S5**).

In contrast, Leger *et al*.^5^ estimate an averaged sunlight to protein production efficiency of between 0.29% (minimum food production efficiency) and 0.87% (maximum feed production efficiency) using a solar PV driven Methanol-RUMP pathway. These results presented here suggest that these efficiencies, at least instantaneously could be pushed almost an order of magnitude higher.

### Conclusion

In this work, we examined a fundamental, molecular-scale model of electromicrobial production of amino acids. It is important to re-state here that this calculation does not predict the performance of any naturally-occurring organism. It simply considers a set of redox transformations and enzymatic reactions, and predicts an upper limit efficiency for any natural or synthetic organism using these reactions.

Electromicrobial protein production could address many issues surrounding modern protein production including greenhouse gas emissions^14,51,52^, nitrogen run-off, and land use^13,14,20,53,54^. Recent results by Leger *et al*.^5^ suggest that the solar to protein conversion efficiency of agriculture could be improved by an order of magnitude by combining PV with electromicrobial production technologies.

We examined electromicrobial protein production systems that assimilate N_2_ using a FeMo nitrogenase reaction; assimilate carbon using one of the six known natural CO_2_-fixation pathways (3HP/4HB, rTCA, WL, 4HB, CBB, 3HP) pathways or assimilate formic acid with the artificial formolase pathway; and uptake electrons and energy through H_2_-oxidation or extracellular electron uptake. The costs of N_2_-fixation mean that electromicrobial protein production is likely never to be as efficient as carbohydrate electromicrobial production. But, our results suggest that they could approach it.

The least efficient system (EEU coupled with the 4HB cycle; EEU-4HB) required 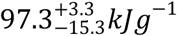 of an average amino acid (**Figure 3B**) (corresponds to an electrical to protein energy conversion efficiency, 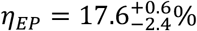; **Figure 3A**). The most efficient system (H_2_-WL) required only 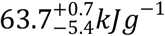 of amino acids (**Figure 3B**) (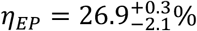, **Figure 3A**). If supplied with electricity by a perfectly efficient single junction Si PV the EEU-4HB system would produce protein with an efficiency of *η_SP_* = 5.8%, while the H_2_-WL system would produce protein with an efficiency of *η_SP_* = 8.9%. These results suggest that the process proposed by Leger *et al*.^5^ could be improved, at least instantaneously, by another order of magnitude.

What’s the best way to achieve the potential of electromicrobial protein production? All of the systems considered in this study rely upon the presence of at least a small amount (≥ a few hundred ppm) O_2_ to generate the maximum amount of reducing equivalents from incoming electrons^25,30,32^. Natural options exist for carbon assimilation in high efficiency engineered EMP systems. For carbon assimilation, the Calvin cycle, 3HP cycle, and Formolase pathway can all be operated in the presence of O_2_. In fact, the H_2_-oxidizing microbe *Ralstonia eutropha* (the chassis organism for the Bionic Leaf which uses the Calvin cycle) fixes CO_2_ in the presence of at least 1% O_2_, while the Fe-oxidizing microbe *Sideroxydans lithotrophicus* ES-1 uses EEU to power CO_2_ fixation in a micro-aerobic environment.

However, N_2_-fixation poses a uniquely formidable challenge for high efficiency electromicrobial production. Over the past decade, several groups have incorporated genes for N_2_-fixation into *E. coli* and demonstrated functional N_2_-fixation^55–60^. But, despite tantalizing possibilities^61^, all known nitrogenase enzymes are sensitive to O_2_. This creates a fundamental incompatibility between EEU and N_2_-fixation that needs to be solved.

Creation of an O_2_-tolerant nitrogenase may be a tall order for evolution. Unlike other enzymes useful in sustainable energy applications like the hydrogenase^62^, there are plenty of evolutionary pressures to drive the creation of an O_2_-tolerant nitrogenase. Despite plenty of demand and opportunities for an O_2_-tolerant nitrogenase to emerge, nature has not presented one.

To date, nature has solved the problem of operating the nitrogenase in an O_2_-rich environment by sequestering it. For example, root nodules in leguminous plants provide an O_2_-shielded environment for symbiotic N_2_-fixing microbes. Likewise filamentous N_2_-fixing cyanobacteria are able to operate the nitrogenase enzyme inside O_2_-impermeable differentiated cells called heterocysts while simultaneously operating oxygenic photosynthesis to generate reducing equivalents in adjacent cells^63^. A similar approach, or recent advances in compartmentalization in synthetic biology^64–68^, give a menu of options for building a synthetic O_2_-resistant compartment for the nitrogenase. Achieving this goal is likely to represent a major challenge in synthetic biology.

Development of an O_2_-resistant compartment will also enable the implementation of highly efficient CO_2_-fixation pathways like the 3HP/4HB cycle, rTCA cycle and Wood-Ljungdahl pathway in synthetic organisms that simultaneously use O_2_ as a metabolic terminal electron acceptor.

Failure to operate enzymatic N_2_-fixation does not spell the end of the road for electromicrobial protein production however. Much as there has been significant development of electrochemical CO_2_ reduction to C_1_ compounds, recent developments in electrochemical N_2_ reduction to ammonia could be a promising complement to biological production of complex amino acids^54^.

## End Notes

### Code Availability

All code used in calculations in this article is available at https://github.com/barstowlab/electrofoods and is archived on Zenodo at https://doi.org/doi:10.5281/zenodo.5698500.

### Materials & Correspondence

Correspondence and material requests should be addressed to B.B.

### Author Contributions

Conceptualization, L.W. and B.B.; Methodology, B.B; Investigation, L.W., S.M., K.R., J.S., M.H., E.N., and B.B; Writing - Original Draft, L.W., S.M., K.R., and B.B.; Writing - Review and Editing, L.W. and B.B.; Resources, B.B.; Supervision, B.B.;

## Acknowledgements

We thank S. Alcaine in the Food Science Department at Cornell University for guidance. This work was supported by Cornell University startup funds (to B.B.), a Burroughs-Wellcome Career Award at the Scientific Interface (to B.B.), and US Department of Energy Biological and Environmental Research award DE-SC0020179. K.R. was supported by a McNair graduate fellowship.

## Competing Interests

The authors declare no competing interests.

## Notes

https://github.com/barstowlab/electrofoods

### Competing Interest Statement

The authors have declared no competing interest.

