## Supplementary Information for "Thermodynamic Constraints on Electromicrobial Protein Production"

#### Supplementary Information Tables

**Table S1.** Symbols used in this article.

**Table S2.** Molecular weights and energy densities for products considered in this article.

**Table S3.** Carbon-fixation and -assimilation, and nitrogen-fixation reactions considered in this article.

**Table S4.** Primary energy inputs for representative protein sources.

**Table S5.** Solar energy costs of photosynthetic protein production.

#### Supplementary Information Datasets

**Dataset S1.** Enzymatic reactions for synthesis of amino acids and D-glucose from metabolic intermediates.

**Dataset S2.** Net molecular input requirements for product synthesis production by 6 naturally-occurring $CO_2$-fixation cycles and the synthetic Formolase formate assimilation pathway.

| Symbol | Unit | Description |
|---|---|---|
| $E_{protein}$ | J molecule$^{-1}$ | Energy carried per amino acid or protein molecule. |
| $\dot{N}_{protein}$ | molecule s$^{-1}$ | Amino acid (or protein) molecules produced per second by electromicrobial production system. |
| $N_A$ | molecule Mol$^{-1}$ | Avogadro constant. |
| F | A s Mol$^{-1}$ | Faraday Constant |
| $P_{e,\ total}$ | J s$^{-1}$ | Total electrical power input into electromicrobial production system. |
| $M_{protein}$ | g Mol $^{-1}$ | Molecular weight of the protein molecule |
| $e$ | A s | Fundamental charge |
| $\nu_{ep}$ | # | Number of electrons needed for synthesis of an amino acid molecule |
| $\Delta U_{cell}$ | V | Potential difference across bio-electrochemical cell |
| $\nu_{e,\ add}$ | # | Number of electrons needed to convert a $C_1$ compound to a protein product |
| $\nu_r$ | # | Number of primary reduction products to make a molecule of final product |
| $\nu_{er}$ | # | Number of electrons to reduce $CO_2$ to a primary reduction product |
| $\nu_{Cr}$ | # | Number of carbon atoms per primary reduction product |
| $\xi_{I2}$ | # | Faradaic efficiency of the bio-electrochemical cell |
| $\xi_{I1}$ | # | Faradaic efficiency of the primary abiotic cell |
| $\xi_C$ | # | Carbon transfer efficiency from cell 1 to cell 2 |
| $\nu_{p,\ NADH}$ | # | Number of NAD(P)H molecules needed to make a final protein molecule |
| $\nu_{p,\ Fd}$ | # | Number of Fd molecules needed to make a final protein molecule |
| $\nu_{p,\ ATP}$ | # | Number of ATP molecules needed to make a final protein molecule |
| $\Delta G_{ATP/ADP}$ | J | Free energy for regeneration of ATP |
| $\Delta U_{membrane}$ | V | Inner membrane potential difference |
| $U_{H2}$ | V | Standard potential of proton reduction to $H_2$ |
| $U_{acceptor}$ | V | Standard potential of terminal electron acceptor reduction |

| Symbol | Unit | Description |
|---|---|---|
| $U_Q$ | V | Redox potential of the inner membrane electron carrier |
| $U_{NADH}$ | V | Standard potential of NADH |
| $U_{Fd}$ | V | Standard potential of Ferredoxin |
| $C_{EP}$ | J $g^{-1}$ | Electrical energy cost per unit mass |
| $C_{SP}$ | ¢ $g^{-1}$ | Financial cost efficiency per unit mass |
| $\eta_{EP}$ | % | Electrical to product (*e.g.,* protein) energy conversion efficiency |
| $\eta_{SP}$ | % | Solar to product (*e.g.,* protein) energy conversion efficiency |

**Table S1.** Symbols used in this article.

| Amino Acid | Symbol | Molecular Weight (Da) | Energy Density (kJ $g^{-1}$) | Energy Density (J $molecule^{-1}$) | Molecular Formula | Carbons per Molecule |
|---|---|---|---|---|---|---|
| Alanine | ALA | 89 | 13.98 | 2.066E-18 | $C_3H_7NO_2$ | 3 |
| Arginine | ARG | 174 | 13.58 | 3.924E-18 | $C_6H_{14}N4O_2$ | 6 |
| Asparagine | ASN | 132 | 9.38 | 2.056E-18 | $C_4H_8N_2O3$ | 4 |
| Aspartate | ASP | 133 | 9.23 | 2.039E-18 | $C_4H_7NO_4$ | 4 |
| Cysteine | CYS | 121 | 17.44 | 3.504E-18 | $C_3H_7NO_2S$ | 3 |
| Glutamine | GLN | 146 | 12.69 | 3.077E-18 | $C_5H_{10}N_2O_3$ | 5 |
| Glutamate | GLU | 147 | 12.53 | 3.059E-18 | $C_5H_9NO_4$ | 5 |
| Glycine | GLY | 75 | 8.67 | 1.080E-18 | $C_2H_5NO_2$ | 2 |
| Histidine | HIS | 155 | 13.57 | 3.493E-18 | $C_6H_9N_3O_2$ | 6 |
| Isoleucine | ILE | 131 | 23.80 | 5.177E-18 | $C_6H_{13}NO_2$ | 6 |
| Leucine | LEU | 131 | 23.80 | 5.177E-18 | $C_6H_{13}NO_2$ | 6 |
| Lysine | LYS | 146 | 20.02 | 4.854E-18 | $C_6H_{14}N_2O_2$ | 6 |
| Methionine | MET | 149 | 21.41 | 5.297E-18 | $C_5H_{11}NO_2S$ | 5 |
| Phenylalanine | PHE | 165 | 25.06 | 6.866E-18 | $C_9H_{11}NO_2$ | 9 |
| Proline | PRO | 115 | 20.10 | 2.039E-18 | $C_5H_9NO_2$ | 5 |
| Serine | SER | 105 | 10.36 | 1.806E-18 | $C_3H_7NO_3$ | 3 |
| Threonine | THR | 119 | 13.94 | 2.755E-18 | $C_4H_9NO_3$ | 4 |
| Tryptophan | TRP | 204 | 23.39 | 7.924E-18 | $C_{11}H_{12}N_2O_2$ | 11 |
| Tyrosine | TYR | 181 | 21.77 | 6.543E-18 | $C_9H_{11}NO_3$ | 9 |
| Valine | VAL | 117 | 21.27 | 4.132E-18 | $C_5H_{11}NO_2$ | 5 |
| Average Amino Acid | AVE | 136.75 | 16.7995 | 3.815E-18 | - | 5 |
| Glucose | GLUC | 180.16 | 15.55 | 4.652E-18 | $C_6H_{12}O_6$ | 6 |
| Butanol | BUT | 74.12 | 36.6 | 4.505E-18 | $C_4H_{10}O$ | 4 |

**Table S2.** Molecular weights and energy densities for products considered in this article. Data for amino acids from May *et al*. [May1990a].

| Reaction | Reference |
|---|---|
| ***1. Calvin Cycle*** | |
| 2 $CO_2$ + 7 ATP + 4 NADH → 1 Acetyl-CoA | Salimijazi *et al*. [Salimijazi2020b]. |
| 3 $CO_2$ + 7 ATP + 5 NADH → 1 Pyruvate | Salimijazi *et al* [Salimijazi2020b]. |
| ***2. Wood-Ljungdahl Pathway*** | |
| 4 $CO_2$ + 2 ATP + 8 NADH → 2 Acetyl-CoA | Berg [Berg2011a]. |
| 2 $Fd_{red}$ + Acetyl-CoA + $CO_2$ → Pyruvate | KEGG R01196. |
| ***3. Reductive TCA Cycle*** | |
| 4 CO2 + 4 ATP + 8 NADH → 2 Acetyl-CoA | Alissandratos *et al.* [Alissandratos2015a], Claassens *et al.* [Claassens2016a]. |
| 2 $Fd_{red}$ + Acetyl-CoA + $CO_2$ → Pyruvate | KEGG R01196. |
| ***4. 3-hydroxypropionate/4-hydroxybutyrate Cycle*** | |
| 6 $HCO_3^-$ + 10 ATP + 10 NADH → 2 pyruvate | Berg *et al*. [BergI2007a], Claassens *et al*. [Claassens2016a]. |
| 2 Pyruvate → 2 Acetyl-CoA + 2 NADH + 2 $CO_2$ | Berg [Berg2002a], Schomburg *et al.* [Schomburg2017a]. |
| ***5. 3-hydroxypropionate Cycle*** | |
| 6 $HCO_3^-$ + 10 ATP + 12 NADH → 2 Pyruvate | Zarzycki *et al*. [Zarzycki2009a], Herter *et al*. [Herter2002a], Berg [Berg2002a]. |
| 2 Pyruvate → 2 Acetyl-CoA + 2 NADH + 2 $CO_2$ | Zarzycki *et al*. [Zarzycki2009a], Herter *et al*. [Herter2002a], Berg [Berg2002a]. |
| ***6. 4-hydroxybutyrate Cycle*** | |
| 1 $CO_2$ + 1 $HCO_3^-$ + 3 ATP + 1 NADH + 6 $Fd_{red}$ → 1 Acetyl-CoA | Huber *et al*. [Huber2008a]. |
| 2 Pyruvate → 2 Acetyl-CoA + 2 NADH + 2 $CO_2$ | Berg [Berg2002a], Schomburg *et al.* [Schomburg2017a]. |
| ***7. Formolase Pathway*** | |
| 6 $HCO_2^-$ + 10 ATP + 4 NADH → 2 3-PG | Siegel *et al*. [Siegel2015a], Bar-Even *et al*. [Bar-Even2016a]. |
| 2 3-PG → 2 Pyruvate + 2 ATP | Berg [Berg2002a]. |
| 2 Pyruvate → 2 Acetyl-CoA + 2 NADH + 2 $CO_2$ | Berg [Berg2002a], Schomburg *et al.* [Schomburg2017a]. |

**Table S3.** $CO_2$-fixation and $C_1$-assimilation reactions. $CO_2$-fixation and $C_1$-assimilation reactions considered in this article were first assembled in Salimijazi *et al*. [Salimijazi2020b] and are restated here for convenience. Overall reactions for production of metabolic intermediates by 6 naturally-occurring $CO_2$-fixation cycles and the synthetic Formolase formate assimilation pathway, and the FeMoCo nitrogenase $N_2$-fixation reaction. Reactions can be referenced KEGG database [Kanehisa2000a, Kanehisa2019a, Kanehisa2021a]. $Fd_{red}$: Reduced Ferredoxin; 3-PG: 3-Phosphoglycerate.

| Food Source | Energy Consumption (MJ $kg^{-1}$ protein) | References | Notes |
|---|---|---|---|
| Beef | 187 - 273 | Williams *et al.* [Williams2006a] | All selected LCAs use a "Cradle to Farm/Factory-Gate" strategy for assessing energy use. This means no transport, retail or household considerations for the most apples-to-apples comparison possible. These numbers are quite variable depending on the inputs per LCA. |
| Pork | 119 - 129 | Williams *et al.* [Williams2006a] | |
| Chicken | 80 - 96 | Williams *et al.* [Williams2006a] | |
| Eggs | 87 - 95 | Williams *et al.* [Williams2006a] | |
| Soybeans | 44.22 | Pimentel *et al.* [Pimentel2009a] | Solar energy input is not considered. |
| Milk/Dairy | 67 - 68 | Williams *et al.* [Williams2006a] | |
| Insect | 170 | Oonincx *et al.* [Oonincx2012a] | This is an example where MJ $kg^{-1}$ food and MJ $kg^{-1}$ protein makes a huge difference. |
| Cultured Meat | 131.9 - 166.5 | Tuomisto *et al.* [Tuomisto2011a] | Divided by protein content of 19.1% as per reference (similar to lean meat protein %) |

**Table S4.** Primary energy inputs for representative protein sources.

| Crop | Country | Protein Produced (g $m^{-2}$ $yr^{-1}$) | Energy Stored in Protein (kJ $m^{-2}$ $yr^{-1}$) | Median Irradiance kJ $m^{-2}$ $yr^{-1}$ | Solar Energy Cost, $C_{SP}$ (kJ $g^{-1}$) | Solar to Protein Energy Conversion Efficiency, $\eta_{SP}$ (%) | Solar to Protein Energy Conversion Efficiency Error (%) |
|---|---|---|---|---|---|---|---|
| Maize | USA | 93 | 1,556 | 5,684,400 | 61,123 | 0.027% | 0.003% |
| Maize | Bulgaria | 63 | 1,050 | 5,061,600 | 80,343 | 0.021% | 0.005% |
| Maize | India | 22 | 372 | 8,967,600 | 407,618 | 0.004% | 0.001% |
| Maize irrigated | USA | 118 | 1,975 | 6,130,800 | 51,956 | 0.032% | 0.002% |
| Maize sillage | USA | 73 | 1,218 | 5,194,800 | 71,162 | 0.023% | 0.003% |
| Sugar beet | USA | 99 | 1,648 | 5,400,000 | 54,545 | 0.030% | 0.003% |
| Soybean | USA | 120 | 2,004 | 5,677,200 | 47,310 | 0.035% | 0.005% |

**Table S5.** Solar energy inputs for protein production by crops. Adapted from Table S1G in Leger *et al*. [Leger2021a]. Solar energy costs per gram of protein were added by us, and units were converted to match those in our article.

### Supplementary Information References


[Alissandratos2015a] A. Alissandratos and C. J. Easton. "Biocatalysis for the application of $CO_2$ as a chemical feedstock". *Beilstein Journal of Organic Chemistry* 11 (2015), pp. 2370–2387. doi:10.3762/bjoc.11.259.

[Bar-Even2016a] A. Bar-Even. "Formate Assimilation: The Metabolic Architecture of Natural and Synthetic Pathways." *Biochemistry* 55 (2016), pp. 3851–63. doi:10.1021/acs.biochem.6b00495.

[Barstow2021a] B. Barstow. Electrofoods. doi:10.5281/zenodo.5698500.

[Berg2002a] J. Berg, J. Tymoczko, and L. Stryer. Biochemistry. 5th. New York, NY: W H Freeman, 2002.

[Berg2011a] I. A. Berg. "Ecological Aspects of the Distribution of Different Autotrophic $CO_2$ Fixation Pathways". *Applied and Environmental Microbiology* 77 (2011), pp. 1925–1936. doi:10.1128/aem.02473-10.

[BergI2007a] I. A. Berg, D. Kockelkorn, W. Buckel, and G. Fuchs. "A 3-Hydroxypropionate/4-Hydroxybutyrate Autotrophic Carbon Dioxide Assimilation Pathway in Archaea". *Science* 318 (2007), pp. 1782–1786. doi:10.1126/science.1149976.

[Claassens2016a] N. J. Claassens, D. Z. Sousa, V. A. M. dos Santos, W. M. de Vos, and J. van der Oost. "Harnessing the power of microbial autotrophy". *Nature Reviews Microbiology* 14 (2016), pp. 692–706. doi:10.1038/nrmicro.2016.130.

[Herter2002a] S. Herter, G. Fuchs, A. Bacher, and W. Eisenreich. "A bicyclic autotrophic $CO_2$ fixation pathway in *Chloroflexus aurantiacus*". *Journal of Biological Chemistry* 277 (2002), pp. 20277–20283. doi:10.1074/jbc.m201030200.

[Huber2008a] H. Huber, M. Gallenberger, U. Jahn, E. Eylert, I. A. Berg, D. Kockelkorn, W. Eisenreich, and G. Fuchs. "A dicarboxylate/4-hydroxybutyrate autotrophic carbon assimilation cycle in the hyperthermophilic *Archaeum Ignicoccus hospitalis*". *Proceedings of the National Academy of Sciences* 105 (2008), pp. 7851–7856. doi:10.1073/pnas.0801043105.

[Kanehisa2000a] M. Kanehisa and S. Goto. "KEGG: Kyoto Encyclopedia of Genes and Genomes". *Nucleic Acids Research* 28.1 (2000), pp. 27–30. doi:10.1093/nar/28.1.27.

[Kanehisa2019a] M. Kanehisa. "Toward understanding the origin and evolution of cellular organisms". Protein *Science* 28.11 (2019), pp. 1947–1951. doi:10.1002/pro.3715.

[Kanehisa2021a] M. Kanehisa, M. Furumichi, Y. Sato, M. Ishiguro-Watanabe,and M. Tanabe. "KEGG: integrating viruses and cellular organisms". *Nucleic Acids Research* 49.D1 (2020), gkaa970–. doi:10.1093/nar/gkaa970.

[Leger2021a] D. Leger, S. Matassa, E. Noor, A. Shepon, R. Milo, and A. Bar-Even. "Photovoltaic-driven microbial protein production can use land and sunlight more efficiently than conventional crops". *Proceedings of the National Academy of Sciences* 118.26 (2021), e2015025118. doi:10.1073/pnas.2015025118.

[May1990a] M.E. May, and J.O. Hill. "Energy content of diets of variable amino acid composition". *Am J Clin Nutrition* 52, 770–776 (1990). doi:10.1093/ajcn/52.5.770.

[Pimentel2009a] D. Pimentel. "Energy Inputs in Food Crop Production in Developing and Developed Nations". *Energies* 2.1 (2009), pp. 1–24. doi:10.3390/en20100001.

[Salimijazi2020b] F. Salimijazi, J. Kim, A. M. Schmitz, R. Grenville, A. Bocarsly, and B. Barstow. "Constraints on the Efficiency of Engineered Electromicrobial Production". *Joule* 4 (2020), pp. 2101–2130. doi:10.1016/j.joule.2020.08.010.

[Schomburg2017a] I. Schomburg, L. Jeske, M. Ulbrich, S. Placzek, A. Chang,and D. Schomburg. "The BRENDA enzyme information system–From a database to an expert system". *Journal of Biotechnology* 261 (2017), pp. 194–206. doi:10.1016/j.jbiotec.2017.04.020.

[Siegel2015a] J. B. Siegel, A. L. Smith, S. Poust, A. J. Wargacki, A.Bar-Even, C. Louw, B. W. Shen, C. B. Eiben, H. M.Tran, E. Noor, J. L. Gallaher, J. Bale, Y. Yoshikuni, M. H. Gelb, J. D. Keasling, B. L. Stoddard, M. E. Lidstrom, and D. Baker. "Computational protein design enables a novel one-carbon assimilation pathway". *Proceedings of the National Academy of Sciences* 112 (2015), p. 3704-3709. doi:10.1073/pnas.1500545112.

[Tuomisto2011a] H. L. Tuomisto and M. J. T. d. Mattos. "Environmental Impacts of Cultured Meat Production". *Environmental Science & Technology* 45.14 (2011), pp. 6117–6123. doi:10.1021/es200130u.

[Williams2006a] A. Williams, E. Audsley, and D. Sandars. "Determining the environmental burdens and resource use in the production of agricultural and horticultural commodities. Defra project report IS0205". Tech. rep. Cranfield University and Defra, 2006.

[Zarzycki2009a] J. Zarzycki, V. Brecht, M. Müller, and G. Fuchs. "Identifying the missing steps of the autotrophic 3-hydroxypropionate $CO_2$ fixation cycle in *Chloroflexus aurantiacus*". *Proceedings of the National Academy of Sciences* 106 (2009), p. 21317. doi.10.1073/pnas.0908356106.
